# Functional correlation between WRN and SAMHD1 in end-resection

**DOI:** 10.1101/2024.02.15.580497

**Authors:** Benedetta Perdichizzi, Pietro Pichierri

## Abstract

Double-strand breaks (DSBs) can cause chromosome rearrangements, leading to cancer and some genetic diseases. WRN and SAMHD1 are proteins implicated in DSB processing and form a complex. Our study shows that SAMHD1 influences the nuclear recruitment of WRN in response to CPT-induced DSBs. Silencing SAMHD1 restores single-stranded DNA formation in WRN-deficient cells. However, DSB accumulation from CPT treatment is not recovered in WRN S1133A or WS cells when SAMHD1 is silenced. This suggests SAMHD1 cooperates with WRN in DNA damage repair and may have additional protective roles when WRN function in DSBs processing is impaired.

## Background, Results and Discussion

The WRN protein is a member of the human RecQ helicases and is found mutated in the genetic disease Werner syndrome (WS) (Hickson, 2003). WRN is a pleotropic protein involved in many DNA metabolisms that has been involved in the processing and repair of DNA double-strand breaks (DSBs) at the DNA replication fork (Mukherjee et al., 2018). DNA double-strand break (DSB) repair by homologous recombination (HR) is initiated by CtIP/MRN-mediated DNA end-resection to maintain genome integrity (Daddacha et al., 2017).

In particular, WRN participates in the long-range stage of the end-resection phase of HR collaborating with the DNA2 nuclease (Palermo et al., 2016; Sturzenegger et al., 2014). SAMHD1 is a dNTP hydrolase that is found mutated in the Aicardi-Goutières syndrome (Beloglazova et al., 2013). SAMHD1 is primary involved in regulating the abundance of the dNTP pool in cells but it is also a crucial player in the native immunity and regulation of inflammation (Sze et al., 2013). More recently, SAMHD1 was found to interact with CtIP and contribute to DNA repair by HR because of the promotion of end-resection at DSBs induced by CPT (Daddacha et al., 2017). DSBs are the most dangerous DNA lesions as they can directly promote chromosome rearrangements if unrepaired or misrepaired so the understanding of the factor contributing to correct repair is critical. Moreover, mutation or deficiency of proteins involved in the DNA damage response (DDR) and DNA repair have been reported to correlate with inflammation. For instance, loss of the WRN has been shown to upregulate inflammatory pathways at the transcriptional level (Turaga et al., 2009). Thus, given that WRN and SAMHD1 both participate in the same pathway of DNA repair and since each deficiency might has an impact on inflammatory or immune response, we focused our attention on the study of whether they could interact and work in a common pathway.

In response to DNA replication stress, WRN is phosphorylated primarily by ATR (Mukherjee et al., 2018). Proteomic analysis of anti-WRN immunoprecipitate from WS cells stably expressing wild-type WRN (WRN^WT^), its ATR-nonphosphorylatable (WRN^3A^), or phosphomimetic (WRN^3D^) mutant provided some new interactors with few of them detected only in the phosphomimetic mutant (see Table in Figure 1a). Interestingly, SAMHD1 was found among the interactors of WRN^3D^ in response to HU (Figure 1a), suggesting that association between WRN and SAMHD1 occurs downstream DNA replication stress or DNA damage after phosphorylation of WRN.

**Figure 1.**
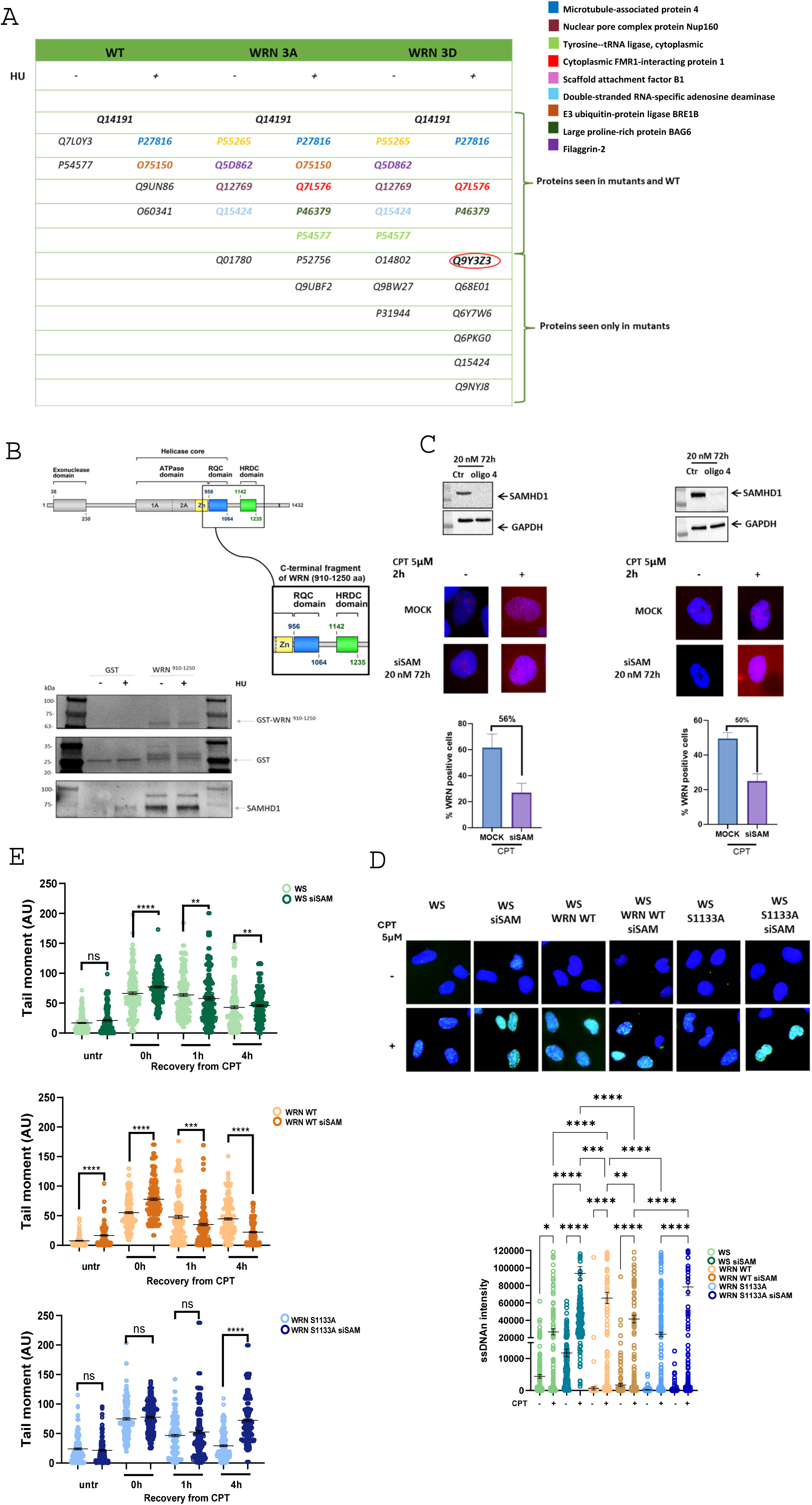
SAMHD1 silencing affects WRN nuclear localization and end resection. **A**. In the table, some of the interactors of WRN WT, WRN 3A, and WRN 3D are reported under conditions of absence and presence of HU. Highlighted in red is the SAMHD1 protein, which from these proteomic data appears to be an interactor of the WRN 3D mutant. **B**. Top: representation of the domains of the WRN protein with focus on the C-terminal fragment from amino acids 910 to 1250. Bottom: pull-down of nuclear extracts from HEK293T cells treated with 2 mM HU for 4 hours and incubated with GST or GST-C-terminal WRN fragment. The WRN fragment and SAMHD1 were detected using rabbit anti-GST (Abgent) and anti-SAMHD1 (Bethyl) antibodies, respectively. **C**. On the right, U2OS cells and on the left, RPE1 hTERT cells were transfected with SAMHD1 siRNA and, after the indicated post-transfection time, exposed to 5 μM CPT for 90 minutes. The graph shows the percentage of cells positive for focal WRN staining. **D**. WS fibroblasts were transiently transfected with the indicated WRN-expressing plasmid. WB shows WRN expression levels 48hrs after transfection using anti-WRN antibody. The ssDNA was analysed at different time points, as indicated. The dot plots show the mean intensity of IdU/ssDNA staining for single nuclei (n=300). Data are presented as mean±SE. Representative images of IdU/ssDNA-stained from CPT-treated cells are shown. Statistical analysis was performed by the ANOVA test (ns = not significant; ^*^= P>0.05; ^**^= P < 0.01; ^***^= P < 0.001; ^****^= P < 0.0001). Where not indicated, pairs are not significant. **E**. DSBs repair efficiency analysis. WS-derived SV40-transformed fibroblasts stably expressed Flag-tagged WRN mutants were transfected with SAMHD1 siRNA and, after the indicated post-transfection time, exposed to 5 μM CPT for 90 minutes and then released in drug free medium at different time points. DSBs repair was evaluated by the neutral Comet assay. In the graph, data are presented as mean tail moment±SE. Representative images from the neutral Comet assay are shown. Statistical analysis was performed by the Student’s t-test (ns = not significant; ^**^= P < 0.001; ^***^P< 0.01; ^****^= P < 0.001).

To confirm interaction between WRN and SAMHD1 and give preliminary data on the region of WRN involved, we performed a GST pull-down assay using the C-terminal region of WRN as bait. Indeed, almost all the protein-protein interactions of WRN are reported to involve the RQC and HRDC domains (Cheng & Bohr, 2003). Thus, we generated a fragment containing both domains (Figure 1b) to pull-down proteins from cell extracts. The first results we obtained indicated that SAMHD1 interacts with the WRN fragment containing the region ranging from amino acid 910 to amino acid 1250 (Figure 1b). Pull-down with HU-treated HEK293T cell extracts revealed no differences in the interaction between WRN^910-1250^ and SAMHD1. Further experiments with purified proteins are needed to confirm a direct interaction between WRN and SAMHD1, via the C-terminal domain, and the stimulation by DNA damage or replication stress in the cell.

Previous studies have suggested that SAMHD1 colocalizes with DNA repair foci in cells exposed to genotoxic agents, suggesting that it may play a more direct role in repairing DNA lesions or stalled replication forks (Coggins et al., 2020). Fork stalling occurs when cells are exposed to genotoxic agents such as HU and camptothecin (CPT) or when they encounter sequences that are intrinsically difficult to replicate (Bakhoum et al., 2018).

Since our data showed an interaction between WRN and SAMHD1, and since both are involved in the repair of CPT-induced DSBs, we wanted to investigate whether the nuclear recruitment of WRN was affected by the absence of SAMHD1 (Figure 1c). We analyzed the formation of WRN nuclear foci by immunofluorescence in two cell lines, U2OS and RPE1 hTERT, after treatment with 5μM CPT for 90 minutes. U2OS is an osteosarcoma cell line, one of the most represented tumors in WS patients, while RPE1 hTERT is a normal retinal epithelial cell line. In the absence of treatment and in agreement with previous studies (Ammazzalorso et al., 2010), WRN was localized only in nucleoli but was barely detectable in nuclear foci. After treatment, we observed an increase in WRN foci-positive cells in both U2OS and RPE1 hTERT. Silencing of SAMHD1 in this condition resulted in significantly reduced recruitment of WRN at the nuclear level, although this was not completely abrogated and did not affect the quality of focal staining, at least in U2OS cells (Figure 1c). This demonstrated that the absence of SAMHD1 affects the nuclear recruitment of WRN. Moreover, this result is consistent with the SAMHD1 interaction with CtIP, which is an event occurring upstream the role of WRN in long-range end-resection (Palermo et al.,2016).

To determine whether SAMHD1 and WRN might act in a common pathway in response to CPT-induced DSBs, we evaluated if silencing of SAMHD1 resulted in changes in DNA end-resection. To evaluate end-resection we assessed ssDNA formation using the native IdU/ssDNA assay (Palermo et al., 2023) (Figure 1d). Formation of ssDNA was evaluated in WS cells stably expressing wild-type WRN (WRN^WT^) and, as a control, an end-resection-defective WRN mutant (WRN^S1133A^). The analysis was performed at 90 min of CPT treatment, when ssDNA formation depends on S1133 phosphorylation of WRN (Palermo et al., 2016).

As expected, a reduced end-resection was observed in WS cells, which are devoid of WRN. Strikingly, depletion of SAMHD1 in WS cells rescued ssDNA formation after CPT treatment (Figure 1d). In the WS^WRN WT^ cell expressing wild-type WRN, the silencing of SAMHD1 reduced the end-resection as evaluated by exposure of ssDNA after CPT treatment (Figure 1d), and this is consistent with previous data (Daddacha et al., 2017). In contrast, the silencing of SAMHD1 in the WRN ^S1133A^ mutant did not result in an end-resection defect, as the levels of ssDNA are increased compared to the non-silenced condition (Figure 1d). These results apparently indicate that loss of SAMHD1 might rescue the end-resection defect conferred by loss of WRN or that the ssDNA exposed in the double mutant derives from other events. Further experiments will be needed to define this point. Finally, we evaluated the accumulation of double-strand breaks in response to SAMHD1 silencing by performing a Neutral Comet assay. The analysis was conducted at 90 minutes of CPT treatment, followed by 0, 1 or 4 hours of recovery, when most of the CPT-induced DSBs are repaired (Palermo et al., 2016).

The analysis showed that SAMHD1 silencing resulted in elevated number of DSBs after CPT treatment in all the genetic backgrounds (Figure 1e). In cells expressing WRN wild-type, SAMHD1 silencing resulted in a faster DSBs repair as evidence by the reduction in the Tail moment values (Figure 1e). This result may be consistent with a DSBs repair pathway switch induced by the defect in end-resection that engage a faster repair pathway as the non-homologous end-joining. In contrast, silencing of SAMHD1 in WS cells or in cells expressing the WRN mutant conferring a defect in the long-range end-resection (WRN S1133A) induced a milder improvement in the extinction of DSBs or a defective DSBs repair as apparent at 4gh of recovery in the WRN ^S1133A^ mutant (Figure 1e; bottom). Indeed, The WRN ^S1133A^ mutant, which presents an end resection defect per se, under SAMHD1 silencing conditions, did not seem to recover from the accumulation of DSBs at 4 hours consistent with end-resection independent roles of SAMHD1 as also shown by the data on ssDNA accumulation (see panel d).

## Methods

### Cell lines and culture conditions

The cells used include RPE1 hTert, retinal pigment epithelial cells immortalized with hTert, U2OS osteosarcoma cells, and the U2OS WRN KO cells generated by Crispr/Cas9-mediated editing (generously provided by Dr. Philippe Frit from the University of Toulouse in Toulouse, France). All the cell lines were maintained in Dulbecco’s modified Eagle’s medium (DMEM; Life Technologies) supplemented with 10% FBS 103 (Boehringer Mannheim) and incubated at 37 °C in a humidified 5% CO2 atmosphere. All cell lines have been tested for the presence of mycoplasma by PCR and DAPI-IF.

The SV40-transformed WRN-deficient fibroblast cell line (AG11395-WS) was obtained from Coriell Cell Repositories (Camden, NJ, USA). To produce stable cell lines, AG11395 (WS) fibroblasts were transduced with lentiviruses expressing the full-length cDNA encoding wildtype WRN (WRNWT), S1133A-WRN (WRN^S1133A^), 3AATM-WRN (WRN^3AATM^), or 3DATM-WRN (WTN^3DATM^). HEK293T cells were from American Type Culture Collection.

### Chemicals

Hydroxyurea (HU 98% powder, Sigma-Aldrich) was dissolved in ddH20 and used at 2mM. Camptothecin (Enzo Life Sciences) was dissolved in DMSO and used at 5 μM. Iododeoxyuridine (IdU) (Sigma-Aldrich) was dissolved in sterile DMEM to obtain stock solutions of 2.5 mM and 200 mM and stored at −20 °C.

### Transfection

SAMHD1 siRNA (Dharmacon) was transfected at final concentration of 20 nM. Transfection was performed using INTERFERin (Polyplus) 48/60 h before to perform experiments.

### GST pulldown Assay

The GST and C-terminal fragment of WRN were incubated with 300 ng of 293T cell extracts, treated and untreated with HU. After 16 hours of incubation, the fragments were separated from the beads, and the interaction of SAMHD1 with WRN fragments was measured through densitometric analysis by Western blotting using rabbit anti-GST (Calbiochem), rabbit anti-WRN (Abgent), and rabbit anti-SAMHD1 (Bethyl) antibodies.

### Neutral comet assay

After treatment, cells were embedded in low-melting agarose and spread onto glass slides. After an electrophoretic run of 20’ (6-7 A, 20V), cells were fixed with methanol. DNA was stained with 0.1% GelRed (Biotium) and examined at 20× magnification with an Olympus fluorescence microscope. Slides were analysed with a computerized image analysis system (CometScore, Tritek Corp.). To assess the amount of DNA DSB breaks, computer generated tail moment values (tail length × fraction of total DNA in the tail) were used. Apoptotic cells (smaller comet head and extremely larger comet tail) were excluded from the analysis to avoid 36 artificial enhancement of the tail moment. A minimum of 150 cells were analyzed for each experiment.

### Immunofluorescence assay

Cells were cultured onto 22×22 coverslip in 35mm dishes. To detect WRN intensity the cells were treated with CPT 5 uM. Next, cells were washed with PBS, permeabilized with 0.5% Triton X-100 for 10 min at 4°C and fixed with 2% sucrose, 3%PFA. For WRN intensity detection, cells were incubated with rabbit anti-WRN (Abcam) for 1h at 37°C in 0.1% saponine/BSA in PBS followed by Alexa Fluor 594 79 Anti-Rabbit. Slides were analyzed (at 40x) with Eclipse 80i Nikon Fluorescence 37 Microscope, equipped with a VideoConfocal (ViCo) system. Fluorescence intensity for each sample was then analyzed using ImageJ software.

### Detection of nascent ssDNA IdU assay

To detect nascent ssDNA, cells were labelled for 15 min with 100 μM IdU (Sigma-Aldrich), immediately prior the end of the indicated treatments. For immunofluorescence, cells were washed with PBS, permeabilized with 0.5% Triton X-100 for 10 min at 4 °C and fixed in 3% PFA/2% sucrose. Fixed cells were then incubated with mouse anti-IdU antibody (Becton Dickinson) for 1 h 109 at 37 °C in 1% BSA/PBS, followed by species-specific fluorescein-conjugated secondary antibodies (Alexa Fluor 488 Goat Anti-Mouse IgG (H + L), highly crossadsorbed— Life Technologies). Slides were analysed with Eclipse 80i Nikon Fluorescence Microscope, equipped with a Video Confocal (ViCo) system. For each time point, at least 250 nuclei were examined by two independent investigators and foci were scored at 40×. Quantification was carried out using the ImageJ software. Only nuclei showing >10 bright foci were counted as positive.

## Acknowledgements

This work was supported by Associazione Italiana per la Ricerca sul Cancro (AIRC) to PP (IG n. 21428).

## Authors’ contribution

B.P. performed experiments to analyse WRN interaction, end-resection, protein relocalisation and analysis of DNA damage. B.P. and P.P. analysed data and and wrote the paper. All authors approved the paper.

## Conflict of interest

The authors declare that they do not have any conflict of interest.

## Notes

### Competing Interest Statement

The authors have declared no competing interest.

## References

1. Ammazzalorso, F., Pirzio, L. M., Bignami, M., Franchitto, A., & Pichierri, P. (2010). ATR and ATM differently regulate WRN to prevent DSBs at stalled replication forks and promote replication fork recovery. EMBO Journal, 29(18), 3156–3169. 10.1038/emboj.2010.205

2. Bakhoum, S. F., Ngo, B., Laughney, A. M., Cavallo, J. A., Murphy, C. J., Ly, P., … Cantley, L. C. (2018). Chromosomal instability drives metastasis through a cytosolic DNA response. Nature, 553(7689), 467–472. 10.1038/nature25432

3. Beloglazova, Natalia et al. “Nuclease activity of the human SAMHD1 protein implicated in the Aicardi-Goutieres syndrome and HIV-1 restriction.” The Journal of biological chemistry vol. 288,12 (2013): 8101–8110. 10.1074/jbc.M112.431148

4. Cheng, W. H., & Bohr, V. A. (2003). Diverse dealings of the Werner helicase/nuclease. Science of aging knowledge environment : SAGE KE, 2003(31), PE22. 10.1126/sageke.2003.31.pe22

5. Coggins, S. A., Mahboubi, B., Schinazi, R. F., & Kim, B. (2020). SAMHD1 functions and human diseases. In Viruses (Vol. 12, Issue 4). MDPI AG. 10.3390/v12040382

6. Daddacha, W., Koyen, A. E., Bastien, A. J., Head, P. S. E., Dhere, V. R., … Yu, D. S. (2017). SAMHD1 Promotes DNA End Resection to Facilitate DNA Repair by Homologous Recombination. Cell Reports, 20(8), 1921–1935. 10.1016/j.celrep.2017.08.008

7. Hickson, I. D. (2003). RecQ helicases: Caretakers of the genome. In Nature Reviews Cancer (Vol. 3, Issue 3, pp. 169–178). 10.1038/nrc1012

8. Mukherjee, S., Sinha, D., Bhattacharya, S., Srinivasan, K., Abdisalaam, S., & Asaithamby, A. (2018). Werner syndrome protein and dna replication. In International Journal of Molecular Sciences (Vol. 19, Issue 11). MDPI AG. 10.3390/ijms19113442

9. Palermo, V., Rinalducci, S., Sanchez, M., …, Franchitto, A., & Pichierri, P. (2016). CDK1 phosphorylates WRN at collapsed replication forks. Nature Communications, 7. 10.1038/ncomms12880

10. Palermo V., Malacaria E., Semproni M., Perdichizzi B., …., Franchitto A., Pichierri P., (2023) Switch-Like Phosphorylation of WRN Integrates End-Resection with RAD51 Metabolism at Collapsed Replication Forks, BioRxiv. doi: 10.1101/403808

11. Sturzenegger, A., Burdova, K., Kanagaraj, R., …, Cejka, P., & Janscak, P. (2014). DNA2 cooperates with the WRN and BLM RecQ helicases to mediate long-range DNA end resection in human cells. Journal of Biological Chemistry, 289(39), 27314–27326. 10.1074/jbc.M114.578823

12. Sze, A., Olagnier, D., Lin, R., van Grevenynghe, J., & Hiscott, J. (2013). SAMHD1 host restriction factor: a link with innate immune sensing of retrovirus infection. Journal of molecular biology, 425(24), 4981–4994. 10.1016/j.jmb.2013.10.022

13. Turaga, R. V. N., Paquet, E. R., Sild, M., Vignard, J., …, Masson, J. Y., & Lebel, M. (2009). The Werner syndrome protein affects the expression of genes involved in adipogenesis and inflammation in addition to cell cycle and DNA damage responses. 10.4161/cc.8.13.8925

